# Enhancing role of nitrogen fixation in biogeochemical cycles of the Pacific Arctic

**DOI:** 10.1101/2025.03.15.643424

**Authors:** Takuhei Shiozaki, Amane Fujiwara, Eiji Watanabe, Shigeto Nishino, Naomi Harada, Akiko Makabe

**Affiliations:** Atmosphere and Ocean Research Institute, The University of Tokyo, Chiba, Japan; Institute of Arctic Climate and Environment Research, Japan Agency for Marine-Earth Science and Technology, Kanagawa, Japan; Institute for Extra-cutting-edge Science and Technology Avant-garde Research, Japan Agency for Marine-Earth Science and Technology, Yokosuka, Japan

## Abstract

The Arctic Ocean is warming rapidly, prompting increased attention to the ecosystem’s response to sea-ice loss and the influx of organisms from lower latitudes. Here we found that nitrogen fixation in the Pacific Arctic region was significantly elevated when sea ice in the Chukchi shelf region retreated earlier. UCYN-A2 was determined to be the dominant diazotroph, with numerical simulation and water mass analyses implying its origin to be in the Bering Sea. UCYN-A2 abundance was positively correlated with temperature, and it extended into the Arctic off-shelf region, which is undergoing oligotrophication as a result of sea-ice loss. In this region, nitrogen fixation contributed significantly to new production (4.6– 100%, average: 34%), underscoring its impact on the biogeochemical cycles. This study highlights the increasing importance of nitrogen fixation in the Pacific Arctic region, driven by warming, sea-ice loss, and borealization.

## Introduction

Arctic surface temperatures are warming at over twice the global average rate^1–3^, with the Arctic Ocean itself experiencing a similar rate of warming in recent years, a phenomenon known as Arctic Ocean Amplification^4^. This phenomenon is attributed to ice–albedo feedback^5^, in conjunction with the increasing inflow of warm water from rivers^6^ and ocean currents from lower latitudes^4,7^. The increasing heat content in the Arctic Ocean is potentially driving a substantial reduction in sea ice^8–10^, with significant implications for the Arctic Ocean ecosystem. Among the most notable impacts is increased primary production across the region^11,12^, attributed to improvements in the underwater light environment and the influx of new nitrogenous nutrients^11^. Therefore, understanding the nitrogen dynamics has become essential^13,14^. Moreover, the Arctic Ocean ecosystem is currently influenced by the migration of subarctic organisms^15–19^, in a phenomenon termed borealization, or more specifically Atlantification and Pacification on the Pacific and Atlantic sides, respectively^20^. If these incoming species alter the ecosystem functions, this effect could also impact biogeochemical cycles. Although this possibility has been suggested in previous studies^15–18^, it has not yet been demonstrated through direct observations.

This study focused on nitrogen fixation in the Arctic Ocean. Nitrogen fixation is a process performed by specific prokaryotes, in which nitrogen gas is converted into ammonia, representing a major nitrogen source in oceans^21^. This process has been studied in tropical and subtropical oligotrophic waters, where it contributes significantly to new production^22–24^. Recent studies have revealed that nitrogen fixation also occurs in polar regions^25–28^. However, its role in the biogeochemical cycles of polar regions has received limited attention, unlike that in tropical and subtropical oligotrophic waters, which have more abundant reactive nitrogen sources, reducing the contribution of nitrogen fixation^13^. In this study, we present evidence that diazotroph influx from the Bering Sea modifies the biogeochemical cycle of the Chukchi and Beaufort Seas. Our results indicate that nitrogen fixation in the Pacific Arctic exhibits considerable interannual variability, corresponding with the distribution of UCYN-A2 carried by water of Pacific origin. Additionally, we demonstrate that the contribution of nitrogen fixation to new production is markedly higher when nitrogen fixation is enhanced.

## Results and Discussion

### High nitrogen fixation and its contribution to new production during late summer

In this study, we examined nitrogen fixation and primary production in the Chukchi and Beaufort Seas on four occasions between 2015 and 2020, from late summer to autumn (Fig. 1). To elucidate the role of nitrogen fixation in the biogeochemical cycle, we simultaneously examined the processes related to new production (nitrate assimilation and nitrification rate). According to oceanographic conventions^29,30^, nitrate-based new production was calculated by subtracting nitrification from the nitrate assimilation rate. The total new production was then estimated by combining the nitrate-based new production with nitrogen fixation. To our knowledge, this is the first study to investigate the interannual and seasonal variability in new production-related processes including nitrogen fixation and nitrification rates across a broad area in this region. In 2015, 2016, and 2017, observations were conducted around the same time in late summer, while in 2020, observations took place in autumn. This variation in observation periods led to differences in average photosynthetic active radiation (PAR) values (Fig. 2a), with significantly lower PAR in 2020 than in other years (*P* < 0.05, Steel–Dwass test). Sea surface temperatures (SSTs) were also markedly lower in 2020 (Fig. 2b). Although average SSTs were similar across the late-summer cruises in 2015, 2016, and 2017, an expansion of high-temperature water masses into the off-shelf region was noted in 2017 (Fig. 3a), a year in which the heat content of Pacific-origin warm water around the Chukchi Borderland reached its highest level in two decades^10^. During the late-summer cruises, surface nitrate concentrations were nearly depleted, except at stations in the Bering Strait (Fig. S1a–c). In contrast, during the autumn cruise, surface nitrate concentrations increased (Fig. 2c), likely due to wind-induced vertical mixing^31^, particularly at stations in the shelf region (Fig. S1d).

**Fig. 1.**
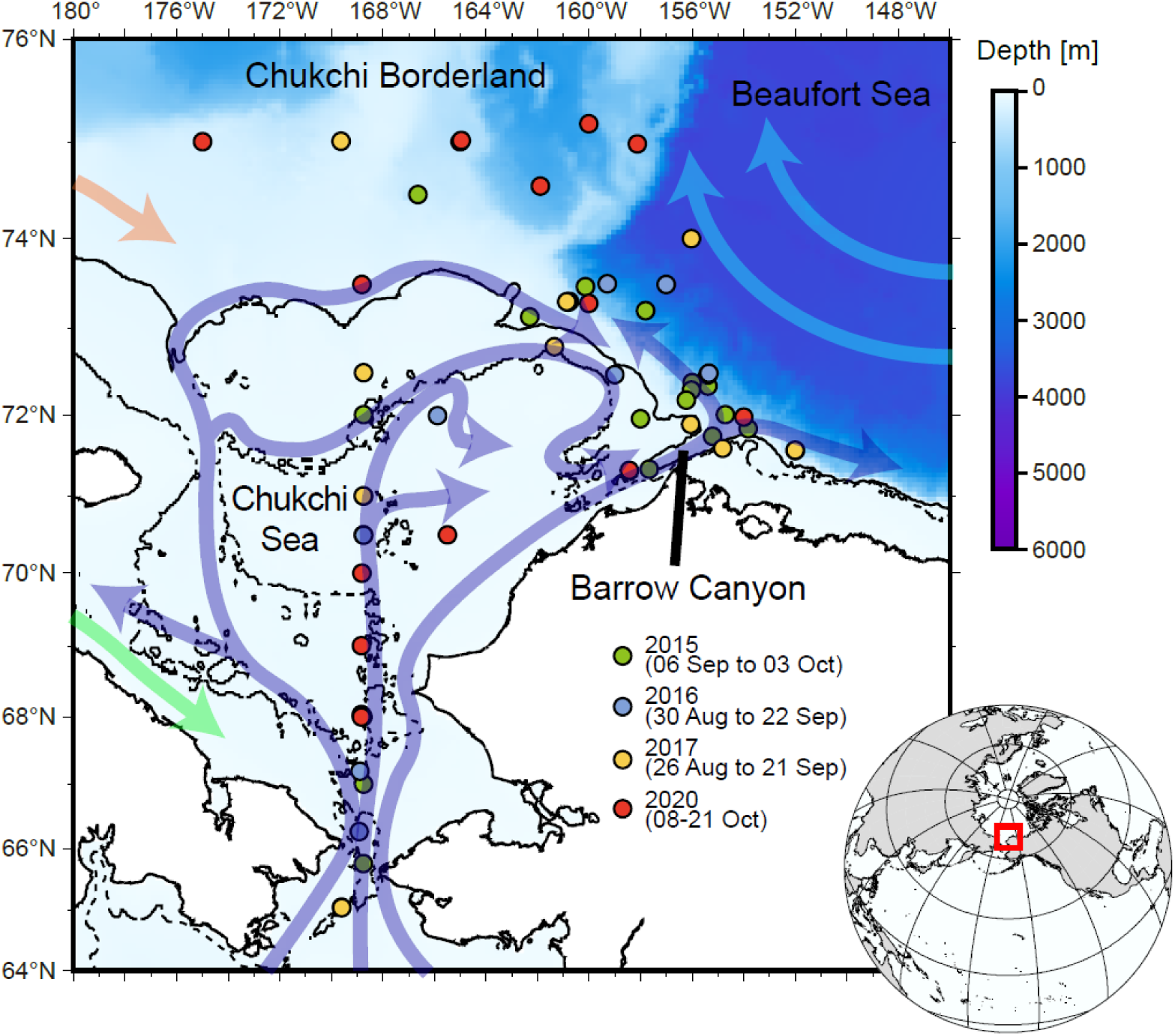
Sampling stations and surface currents in the Chukchi and Beaufort Seas. The current field is based on that of Pickart et al. (2023). Dashed and solid lines indicate 50-m and 100-m isobaths, respectively.

**Fig. 2.**
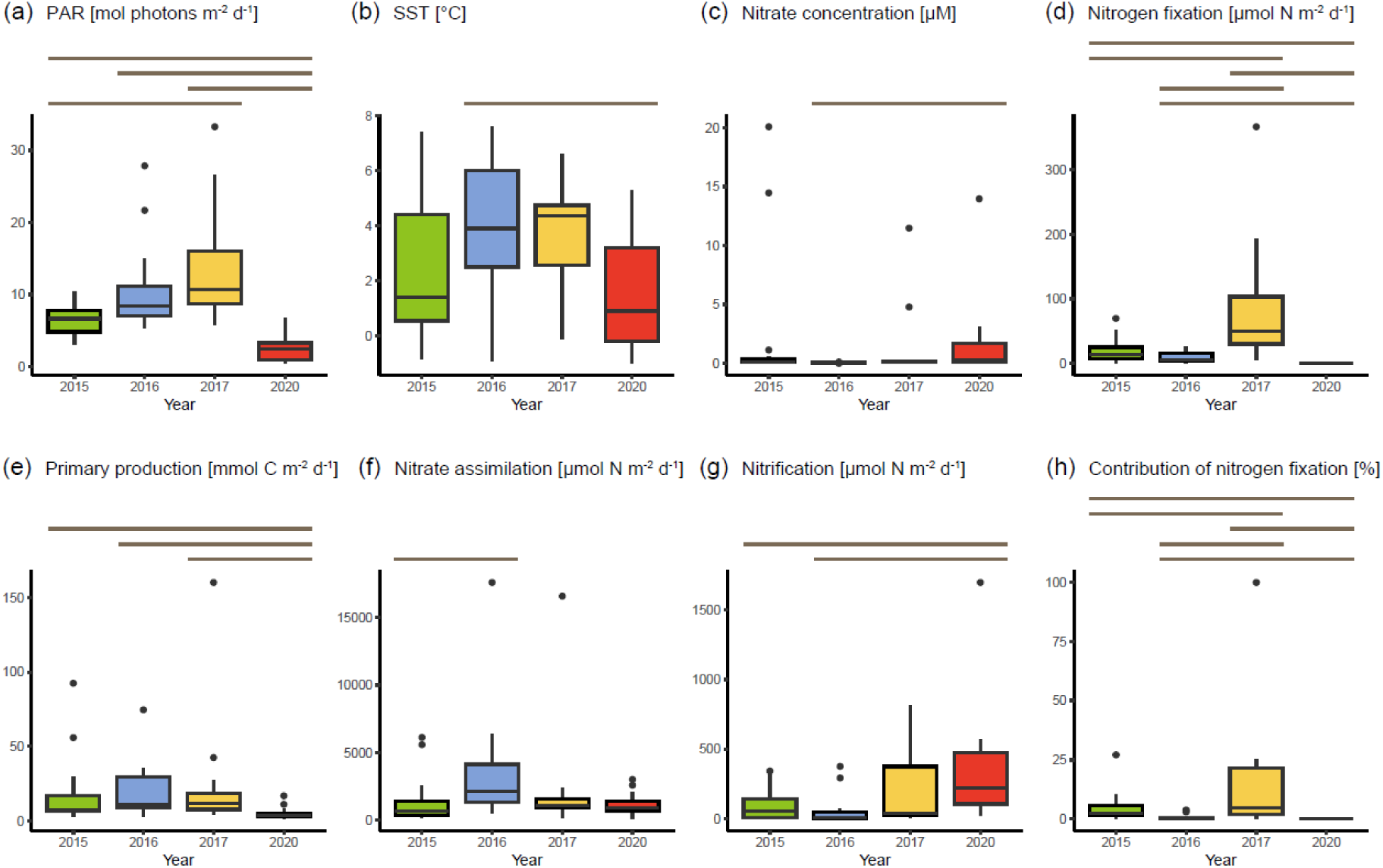
Yearly variation in environmental parameters and biological activity. Box plots show data for (a) photosynthetic active radiation (PAR), (b) sea-surface temperature (SST), (c) surface nitrate concentration, (d) nitrogen fixation, (e) primary production, (f) nitrate assimilation, (g) nitrification, (h) contribution of nitrogen fixation to new production in each year. Lines on box plots indicate significant differences (*P* < 0.05, Steel–Dwass test). For example, there were significant differences in PAR between 2015 and 2017, 2015 and 2020, 2016 and 2020, and 2017 and 2020.

**Fig. 3.**
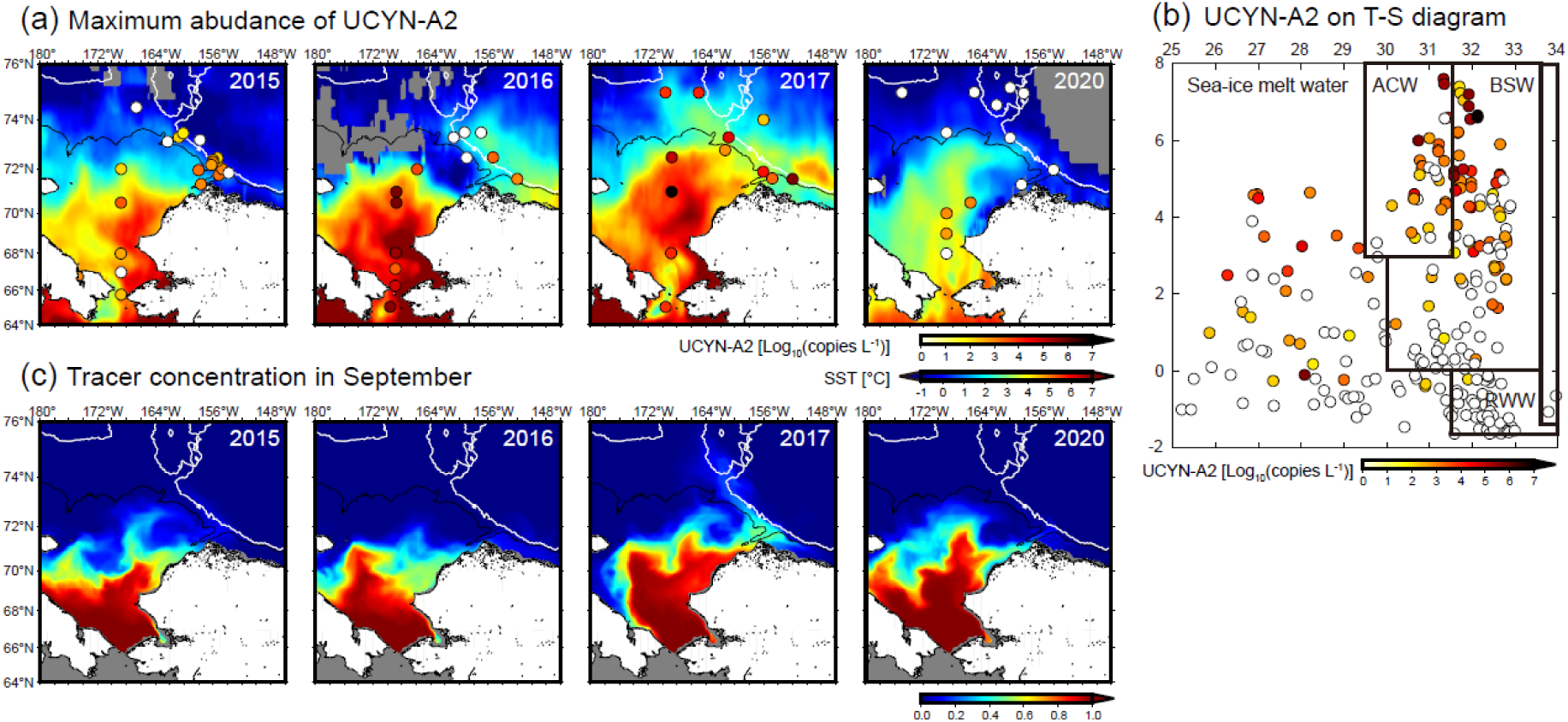
Distribution and potential origin of UCYN-A2 in the Chukchi and Beaufort Seas. (a) Spatial distribution of maximum abundance of UCYN-A2 in each year. Background contours indicate satellite-derived SSTs in September. Gray areas indicate no data. (b) UCYN-A2 abundance on T-S diagram. Water classifications were defined by Pickart et al. (2023). ACW, Alaskan Coastal Water; BSW, Bering Summer Water; RWW, Remnant Winter Water. (c) Simulated distribution of a Pacific water tracer released at the Bering Strait from the time of sea-ice retreat; its concentration in the ocean surface layer in September is shown. Solid black and white lines indicate 100-m and 1000-m isobaths, respectively.

Nitrogen fixation exhibited substantial variation across years and seasons (Fig. 2d). During the late-summer cruises, nitrogen fixation was significantly higher in 2017 than in 2015 and 2016 (*P* < 0.05, Steel–Dwass test), with elevated nitrogen fixation observed not only in the shelf region but also extending to the off-shelf region (Fig. S1). No nitrogen fixation was detected during the autumn cruise. Nitrogen fixation showed no significant correlation with environmental properties such as temperature, salinity, chlorophyll *a* concentration, PAR, or nutrient concentration (Table S1), indicating that *in situ* environmental factors may not be involved in the interannual variation in nitrogen fixation. In contrast to nitrogen fixation, primary production showed no variation among the late-summer cruises; however, it was significantly lower in autumn than in the late-summer cruises (Fig. 2e; *P* < 0.05, Steel–Dwass test). Across all years, primary production showed a significant positive correlation with surface chlorophyll *a* concentrations (Table S1, *P* < 0.001). Nitrate assimilation and nitrification rates also showed no variation among the late-summer cruises (Fig. 2f, g). During these cruises, the nitrate assimilation rate was correlated with surface chlorophyll *a* concentration and depth-integrated primary production (Table S1, *P* < 0.001). In contrast, no significant correlation was observed during the autumn cruise, implying that nitrate assimilation may primarily be carried out by non-primary producers^32^, unlike during the late-summer cruises. Nitrification rates tended to be higher in autumn; thus, as nitrification in this region is inhibited by light^33^, it was likely enhanced during autumn when PAR levels were lower. We calculated the contribution of nitrogen fixation to new production at each station (= nitrogen fixation/[nitrogen fixation + nitrate assimilation – nitrification]). This contribution was generally higher in 2017 (Fig. 2h), a year with notably high nitrogen fixation, particularly in the off-shelf region (17.0–100%), with the exception of one station (St. 74; 4.6%). These high values are comparable to those reported in subtropical oligotrophic oceans^22,24,34^, indicating an increased role of nitrogen fixation in the biogeochemical cycle of off-shelf waters in 2017.

### Nitrogen fixation by UCYN-A2 and its source from the Bering Sea

To examine the factors influencing interannual variation in nitrogen fixation, we further analyzed diazotroph communities in the surface water, where nitrogen fixation was widely detected during the late-summer cruises. Diazotroph community composition varied significantly across the research cruises (Fig. S2). In 2015, ASVs homologous to the *nifH* gene of NB3-*Pelobacter*^35,36^ were dominant at most stations. In 2016, UCYN-A2 was prevalent at most stations in the shelf region; however, non-cyanobacterial diazotrophs, including NB3-*Pelobacter*, PM-RGC_gene_1205376, and AEA49320 were dominant in the off-shelf region. In 2017, UCYN-A2 was the dominant diazotroph at most stations, including the off-shelf region. In 2020, UCYN-A1 and UCYN-A2 were the major diazotrophs at the shelf-region station 7 and the off-shelf station 10, respectively, whereas non-cyanobacterial diazotrophs, including NB3-*Pelobacter*, PM-RGC_gene_1205376, and CE2_5m_1_g, were dominant at other stations. These results imply that UCYN-A2, which was dominant in 2017, may have contributed to the nitrogen fixation anomaly observed that year.

We also quantified the *nifH* gene of UCYN-A2 and detected it in all cruises (Fig. 3a). Notably, UCYN-A2 was widely found at high abundances in the off-shelf regions in 2017, whereas in other cruises, it was primarily confined to the shelf region, except around Barrow Canyon. These distributions were consistent with the diazotroph community composition results. A significant positive correlation was found between nitrogen fixation and UCYN-A2 abundance in 2017 (Fig. S3a, *P* < 0.05), implying that nitrogen fixation was primarily performed by UCYN-A2. As previously reported^37^, UCYN-A2 abundance was significantly positively correlated with water temperature (Fig. S3b, *P* < 0.05). Therefore, the low abundance of UCYN-A2 observed in 2020 was likely attributable to the lower water temperatures during that period. Additionally, low light levels in autumn may have contributed to their reduced abundance. UCYN-A2 was recently found to be associated with photosynthetic organisms as an organelle^38^, implying that the photosynthetic activity of its host declines under low light conditions. These findings imply that it is very difficult for UCYN-A2 to survive during winter in the Arctic Ocean.

Next, we questioned the origin of UCYN-A2 that reached the Arctic Ocean. High UCYN-A2 abundance was detected in water masses identified as the Alaskan Coastal Water and Bering Summer Water^39^, as indicated by a temperature–salinity diagram (Fig. 3b). Therefore, UCYN-A2 likely traveled northward through the Bering Strait into the study region. We further conducted yearly numerical experiments which adopt the Pacific water tracer released at the Bering Strait from the timing of sea-ice retreat. Notably, sea-ice retreat in the Chukchi shelf region occurred significantly earlier in 2017 than in other years (Fig. S4). By September in 2015 and 2016, when the cruise observations were performed, the Pacific water tracer had barely reached the region north of Barrow Canyon (Fig. 3c). However, in 2017, a substantial amount of tracer extended to the Chukchi Borderland region. The distributions of these tracers corresponded closely with the observed distribution of UCYN-A2. These analysis results imply that UCYN-A2 was transported from the Bering Sea into the Arctic Ocean, where it contributed significantly to nitrogen fixation.

### Enhancing role of nitrogen fixation in biogeochemical cycles due to climate change

This study demonstrated that UCYN-A2 transported from subarctic regions actively fixes nitrogen and makes a substantial contribution to new production, particularly in the off-shelf region. Primary production and nitrate-based new production (= nitrate assimilation – nitrification) tended to be lower in this region than in the shelf region, except in 2020 (Fig. S5a, b), due to the significantly deeper nitracline in the off-shelf area compared to the shelf (Fig. S5c). However, nitrogen fixation did not clearly differ between the off-shelf and shelf regions (Fig. S5d). The relative increase in the significance of nitrogen fixation in the deeper nitracline region mirrors a phenomenon commonly observed in tropical and subtropical oligotrophic oceans^24,34,40^.

Previous studies have reported that the basin region of the Beaufort Sea has undergone surface oligotrophication and nitracline deepening, driven by strengthened circulation linked to sea-ice reduction^20,41,42^. While those studies analyzed data up to 2010, the present study extended the analysis period to 2020. Between 2015 and 2020, particularly east of 160°W, nitrate concentrations at a depth of 50 m were found to be extremely low (< 0.5 µM) (Fig. 4a). This oligotrophic area has expanded significantly compared to 2002–2004, when the Beaufort Sea retained extensive summer sea ice. Recent findings indicate that the circulation field has shifted eastward due to ongoing sea-ice reduction^43^, yet the trend in oligotrophication in the basin region persists. Pacific-origin water flowing inward from the Bering Sea primarily reaches the basin area via the Barrow Canyon^44^. Given that UCYN-A2 is likely supplied from the Bering Sea to the Arctic, an earlier sea-ice melting period, such as that observed in 2017, increases the likelihood of UCYN-A2 being transported to the basin region. The timing of sea-ice retreat in the study area has been progressively advancing every year^45^, likely linked to rising heat content. Our satellite data analysis shows a significant increase in surface-water temperatures across the Arctic, including our study area and the oligotrophic Beaufort Sea (Fig. 4b). UCYN-A2 abundance in the Arctic Ocean was significantly positively correlated with temperature, implying that rising water temperatures in the Arctic Ocean promote the growth of host organisms for UCYN-A2. Correctively, the importance of nitrogen fixation in the Arctic Ocean is increasing compared to the period when perennial sea ice extensively covered the region.

**Fig. 4.**
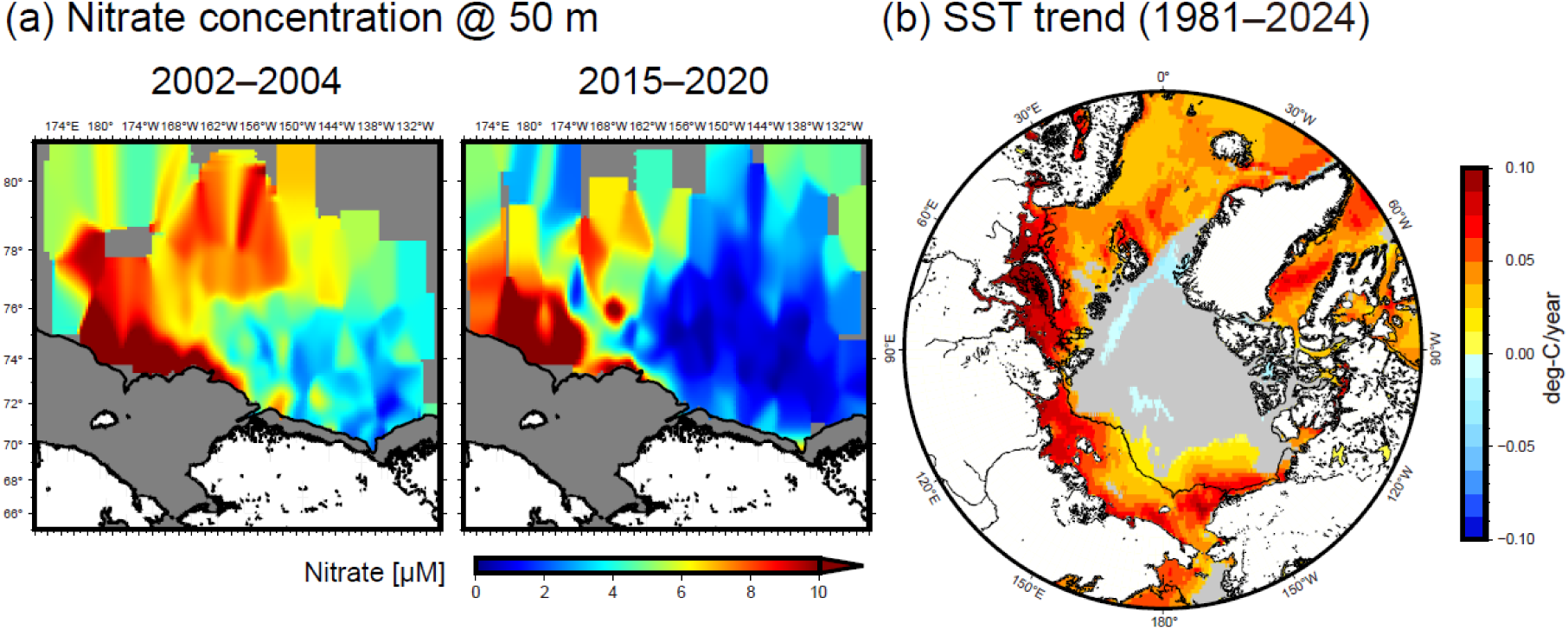
Changes in nutrient fields in the Chukchi and Beaufort Seas and SSTs in the pan-Arctic Ocean. (a) Nitrate concentrations at 50 m during 2002–2004 and 2015–2020. (b) Trends in SSTs during 1981 and 2024. Solid lines indicate 100-m isobaths.

We found that the contribution of nitrogen fixation to new production on the Pacific side of the Arctic Ocean could be increasing due to warming, sea-ice reduction, and pacification. While warming (Fig. 4b) and sea-ice reduction are widespread phenomena across the Arctic^12,46^, environmental conditions on the Atlantic side differ notably from those on the Pacific side. On the Atlantic side, Atlantic water flows directly into the Barents Sea via the Norwegian Sea, where borealization is more pronounced compared to the Pacific side^47^. However, the inflow of Atlantic water and active mixing in the Barents Sea create conditions that facilitate nutrient supply to the surface^20,48^. Additionally, circulation patterns on the Atlantic side are less prone to oligotrophication, unlike conditions observed in the Beaufort Sea^20,49^. Therefore, the phenomena described in this study are unlikely to occur on the Atlantic side. Conversely, it appears that the significant contribution of nitrogen fixation to the Arctic ecosystem is predominantly limited to the Pacific side.

The increasing contribution of nitrogen fixation indicates a significant change in the Arctic nitrogen cycle, but its impact extends beyond nitrogen alone. Nitrogen fixation produces reactive nitrogen from nitrogen gas while simultaneously consuming phosphates. Consequently, in subtropical regions where nitrogen fixation is active, previous studies have reported a decline in phosphate concentrations^50,51^. Notably, in this study, the surface phosphate concentrations (average 0.50 ± 0.01 µM) at stations with a high contribution (≥ 17.0% new production) of nitrogen fixation in the off-shelf region during 2017 were significantly lower compared to the same season in 2015 (0.55 ± 0.05 µM) and 2016 (0.53 ± 0.03 µM) (*P* < 0.05, Wilcoxon rank sum test). This result implies that increased nitrogen fixation alters phosphate concentrations in the Arctic Ocean, thereby influencing the phosphorus cycle. In the Arctic Ocean, the reduction of sea ice has mitigated light limitation, such that nitrogen is the main factor limiting primary production^13^. Consequently, understanding nitrogen dynamics is crucial for comprehending the Arctic ecosystem. The growing significance of nitrogen fixation in this region is poised to reshape our understanding of Arctic biogeochemical cycles. To improve the accuracy of future predictions for the Arctic marine ecosystem, precise modeling of the baseline processes is essential.

## Materials and Methods

Observations and experiments were performed onboard the research vessel (R/V) *Mirai* in 2015 (MR15-03, September 6 to October 3), 2016 (MR16-06, August 30 to September 22), 2017 (MR17-05C, August 26 to September 21), and 2020 (MR20-05C, October 8–21) in the Chukchi and Beaufort Seas. Temperature and salinity profiles were measured using an SBE 911-plus CTD system (Sea-Bird Electronics, Bellevue, WA, USA). The salinity sensor was calibrated using bottle data collected throughout the cruise. Water samples were collected in an acid-cleaned bucket and Niskin-X bottles. In the euphotic zone, water samples were collected from depths corresponding to 100, 10, 1, and 0.1% of PAR, just beneath the surface. The profiles of PAR in the water column were measured using a PRR-800 (2015 and 2017) or C-OPS (2016 and 2020) spectroradiometer (Biospherical Instruments, San Diego, CA, USA) immediately prior to water sampling. We also measured sea surface PAR along the cruise tracks using a shipboard quantum sensor (Li-190R, LI-COR, Lincoln, NE, USA).

### Nutrients and chlorophyll *a*

Nitrate, nitrite, ammonium, and phosphate concentrations were determined on board using the QuAAtro 2-HR system (SEAL Analytical, Southampton, UK). Chlorophyll *a* concentrations were also determined on board using a fluorometer (Turner Designs, San Jose, CA, USA) after extraction with *N,N*-dimethylformamide. The detailed analytical methods of nutrients and chlorophyll *a* are provided in the cruise reports (2015, https://doi.org/10.17596/0001867; 2016, https://doi.org/10.17596/0001870; 2017, https://doi.org/10.17596/0001879; 2020, https://doi.org/10.17596/0002165). The nitracline depth was defined as the depth at which the nitrate concentration became greater than 1 µM.

### Nitrogen fixation and primary production

Nitrogen fixation was determined using the ^15^N_2_ gas dissolution method^52^ for all cruises, and primary production was determined using the ^15^N-^13^C dual inlet technique, following procedures previously reported for this region^27^. Samples for incubation were collected in duplicate in acid-cleaned 1.2-L or 2.3-L polycarbonate bottles. In each cruise, ^15^N_2_ gas produced by SI Science was utilized; ^15^N species contamination from the gas has been confirmed to be negligible^53^. Samples were incubated in an on-deck incubator filled with flowing surface seawater for 24 h. To calculate quantifiable nitrogen fixation rates, we performed a sensitivity analysis of nitrogen fixation measurements^54,55^.

### Nitrate assimilation

Nitrate assimilation was determined by the Michaelis–Menten kinetics approach to correct for overestimation caused by excessive use of the ^15^N tracer^56,57^, with samples collected from the surface and 10% light depth, following procedures previously reported for this region^27^. Samples for incubation were collected in acid-cleaned 1.2-L polycarbonate bottles. Samples were incubated for < 3 h during daytime in an on-deck incubator. Daily nitrate assimilation rates were calculated from daytime and nighttime ratios of nitrate assimilation determined previously for the Chukchi Sea^58^.

### Nitrification

Nitrification rates were also determined using the ^15^N tracer approach^59^ as described in our previous study^33^. Samples for incubation were collected in duplicate in acid-cleaned 0.3-L polycarbonate bottles and incubated for 24 h in an on-deck incubator. The ^15^N/^14^N ratios in nitrate + nitrite of initial and incubated samples were determined using the denitrifier method^60^. The experimental procedures were described in detail elsewhere^33,61^.

### DNA extraction, *nifH* sequencing, and quantitative polymerase chain reaction (PCR)

Samples for DNA analysis (2.3 L) were filtered into Sterivex-GP pressure filter units with a 0.22-µm pore size (Millipore, Billerica, Massachusetts, USA) and stored at –80°C until onshore analysis. Total DNA was extracted using a ChargeSwitch Forensic DNA Purification Kit (Invitrogen, Carlsbad, CA, USA) following the manufacturer’s instructions. Partial *nifH* fragments were amplified using a nested PCR approach with samples collected from the surface, where maximum nitrogen fixation typically occurs. The PCR conditions, polymerase, and product purification methods were consistent with those used in previous studies conducted in the same region^27^.

All samples were sequenced on the MiSeq platform with Reagent Kit v3 (600 cycles) and a Phix control v3 (Illumina, San Diego, CA, USA). Sequenced reads were demultiplexed using MiSeq Reporter v2.6.2 (Illumina). Data analysis was performed with the QIIME2 program v2022.8^62^, after primer sequences were removed using Cutadapt^63^. Reads were denoised and clustered based on sequence variants at single-nucleotide resolution using the DADA2 plug-in^64^. The *nifH* data were translated into amino acid sequences, with non-*nifH* and frameshifted sequences excluded. The resulting sequences were compared against the *nifH* gene catalog of non-cyanobacterial diazotrophs^36^ and the GenBank database using BLASTp.

Quantitative PCR analysis of UCYN-A2 was applied to all DNA samples and performed using the ABI 7500 Real-time PCR system (Applied Biosystems, Foster City, CA, USA) for the 2015, 2016, and 2017 cruises, and the StepOnePlus Real-Time PCR System (Applied Biosystems) for the 2020 cruise. We used previously reported primers and probes for UCYN-A2^65^. The R^2^ values for the standard curves were 0.994–1.000. The efficiency of the qPCR analyses ranged from 94.2 to 101.8%.

### Numerical experiments

Numerical experiments using a virtual tracer were conducted to visualize Pacific water transport from the Bering Strait using the Center for Climate System Research Ocean Component Model (COCO) v4.9^66^. The pan-Arctic regional COCO model used in this analysis has reasonably reproduced major fields of ocean circulation in the western Arctic^67^. The model configuration, experimental design, and corresponding physical hydrographic/circulation results were identical to those reported for previous experiments^43,67^. The horizontal resolution was approximately 5 km, which is sufficient to represent detailed ocean currents such as the Barrow Canyon throughflow. A virtual passive tracer with a concentration value of 1 (assuming Pacific-origin water) was continuously released at the Bering Strait from the time of sea-ice retreat in 2015 (May 22), 2016 (May 16), 2017 (May 13), and 2020 (May 22) (Fig. S4). Tracer values throughout the model domain were maintained at zero until these times in each year. The advection and diffusion of the tracer were calculated along with ocean temperature and salinity.

### Satellite data analyses

Remotely sensed SST data were obtained from the National Oceanic and Atmospheric Administration Optimum (NOAA) Interpolation SST Version 2 product^68^ (http://www.esrl.noaa.gov/psd/data/gridded/data.noaa.oisst.v2.html). The inter-annual SST trend (Fig. 4b) was computed by applying linear regression to each grid. Daily sea-ice concentration (SIC) data derived from Advanced Microwave Scanning Radiometer 2 (AMSR2) were obtained from the Japan Aerospace Exploration Agency (JAXA) web portal (https://gportal.jaxa.jp/gpr/). The timing of sea-ice melt was calculated using daily SIC data for each year. The definition of non-ice-covered pixels was defined at SIC < 20%, and the date of sea-ice retreat was defined as the last date when SIC fell below 20%, prior to the observed annual sea-ice minimum area. In this study, we used SIC data based on a 5-day moving average.

### Historical nutrient data

The R/V *Mirai* data utilized in this study were acquired in 2002, 2004, 2008, 2009, 2010, and 2012–2020. Additionally, we used data obtained from the R/V *Araon* cruise during August 2020 as a part of the “Korea–Arctic Ocean Warming and Response of Ecosystem (K-AWARE)” project. We also used data that have been collected yearly since 2003 by the Canadian Coast Guard Ship *Louis S. St-Laurent* under the collaborative framework of the Beaufort Gyre Exploration Project. This study also included data from the Chukchi Borderland Project^69^ and the International Siberian Shelf Study^70^ conducted by the United States Coast Guard Cutter Polar Star (USA) in 2002 and Yacob Smirniskyi (Russia) in 2008. All data utilized in this study, as well as the observation periods and data sources, are summarized in Table 1.

**Table 1.** Expeditions, observation periods, and data sources.

| Expedition | Observation period | Data source |
| --- | --- | --- |
| *Mirai* 2002 | September 2 – October 10, 2002 | http://www.godac.jamstec.go.jp/darwin/e |
| 2004 | September 1 – October 12, 2004 |  |
| 2008 | August 26 – October 9, 2008 |  |
| 2009 | September 7 – October 15, 2009 |  |
| 2010 | September 2 – October 16, 2010 |  |
| 2012 | September 3 – October 17, 2012 |  |
| 2013 | August 28 – October 21, 2013 |  |
| 2014 | August 31 – October 10, 2014 |  |
| 2015 | August 24 – October 22, 2015 |  |
| 2016 | August 22 – October 5, 2016 |  |
| 2017 | August 23 – October 1, 2017 |  |
| 2018 | October 24 – December 7, 2018 |  |
| 2019 | September 28 – November 10, 2019 |  |
| 2020 | September 19 – November 2, 2020 |  |
| *Araon* 2020 | August 4 – August 31, 2020 | https://kpdc.kopri.re.kr/search/694ee19e-1c0b-4a44-8fd6-e83c94992731 |
| BGEP 2003 | August 7 – September 7, 2003 | https://www2.whoi.edu/site/beaufortgyre/data/ctd-and-geochemistry/ |
| 2004 | July 29 – September 2, 2004 |  |
| 2005 | July 29 – September 1, 2005 |  |
| 2006 | August 5 – September 14, 2006 |  |
| 2007 | July 26 – August 31, 2007 |  |
| 2008 | July 17 – August 21, 2008 |  |
| 2009 | September 17 – October 15, 2009 |  |
| 2010 | September 15 – October 15, 2010 |  |
| 2011 | July 21 – August 18, 2011 |  |
| 2012 | August 2 – September 8, 2012 |  |
| 2013 | August 1 – September 2, 2013 |  |
| 2014 | September 21 – October 17, 2014 |  |
| 2015 | September 20 – October 16, 2015 |  |
| 2016 | September 22 – October 18, 2016 |  |
| 2017 | September 7 – October 2, 2017 |  |
| 2018 | September 7 – October 2, 2018 |  |
| 2019 | September 12 – October 4, 2019 |  |
| 2020 | September 14 – October 2, 2020 |  |
| *CBL* 2002 | August 19 – September 23, 2002 | http://psc.apl.washington.edu/HLD/CBL/CBL.html |
| ISSS 2008 | August 15 – September 26, 2008 | https://cchdo.ucsd.edu/cruise/90JS20080815 |

Expeditions conducted by the R/V *Mirai* (Japan) and R/V *Araon* (Korea) are labeled *Mirai* and *Araon*, respectively, with their expedition years. The Canada/USA Beaufort Gyre Exploration Project using the Canadian Coast Guard Ship *Louis S. St-Laurent* is labeled BGEP. The Chukchi Borderland Project and International Siberian Shelf Study conducted by the US Coast Guard Cutter Polar Star (USA) in 2002 and Yacob Smirniskiy (Russia) in 2008, respectively, are labeled CBL 2002 and ISSS 2008, respectively.

## Acknowledgements

We thank the captain, crew members, and participants of the R/V Mirai Arctic cruises for their cooperation at sea. This study was financially supported by the Japan Society for the Promotion of Science (JSPS) KAKENHI Grant JP23H05411, and JP24K22347, and Arctic Challenge for Sustainability II (ArCS II) of the Ministry of Education, Culture, Sports, Science and Technology.

## References

1 Cohen, J. et al. Recent Arctic amplification and extreme mid-latitude weather. Nature Geoscience 7, 627–637 (2014).

2 Serreze, M. C. & Francis, J. A. The arctic amplification debate. Climatic Change 76, 241–264 (2006).

3 Rantanen, M. et al. The Arctic has warmed nearly four times faster than the globe since 1979. Communications Earth & Environment 3, 168 (2022).

4 Shu, Q. et al. Arctic Ocean Amplification in a warming climate in CMIP6 models. Sci Adv 8, eabn9755 (2022).

5 ACIA. Impacts of a Warming Arctic: Arctic Climate Impact Assessment. ACIA Overview report., (Cambridge University Press, 2004).

6 Park, H. et al. Increasing riverine heat influx triggers Arctic sea ice decline and oceanic and atmospheric warming. Sci Adv 6, eabc4699 (2020).

7 Yamagami, Y., Watanabe, M., Mori, M. & Ono, J. Barents-Kara sea-ice decline attributed to surface warming in the Gulf Stream. Nature communications 13, 3767 (2022).

8 Screen, J. A. & Simmonds, I. The central role of diminishing sea ice in recent Arctic temperature amplification. Nature 464, 1334–1337 (2010).

9 Timmermann, M.-L., Toole, J. & Krishfield, R. Warming of the interior Arctic Ocean linked to sea ice losses at the basin margins. Sci Adv 4, eaat6773 (2018).

10 Muramatsu, M. et al. Subsurface warming associated with Pacific Summer Water transport toward the Chukchi Borderland in the Arctic Ocean. Scientific reports 15, 24 (2025).

11 Lewis, K. M., van Dijken, G. L. & Arrigo, K. R. Changes in phytoplankton concentration now drive increased Arctic Ocean primary production. Science 369, 198–202 (2020).

12 Ardyna, M. & Arrigo, K. R. Phytoplankton dynamics in a changing Arctic Ocean. Nature Climate Change 10, 892–903 (2020).

13 Tremblay, J. E. et al. Global and regional drivers of nutrient supply, primary production and CO2 drawdown in the changing Arctic Ocean. Progress in Oceanography 139, 171–196 (2015).

14 Arrigo, K. R., Mills, M. M. & Juranek, L. W. The Arctic Ocean Nitrogen Cycle. Journal of Geophysical Research: Biogeosciences 129 (2024).

15 Reid, P. C. et al. A biological consequence of reducing Arctic ice cover: arrival of the Pacific diatom Neodenticula seminae in the North Atlantic for the first time in 800,000 years. Global Change Biology 13, 1910–1921 (2007).

16 Kortsch, S., Primicerio, R., Fossheim, M., Dolgov, A. V. & Aschan, M. Climate change alters the structure of arctic marine food webs due to poleward shifts of boreal generalists. Proc. R. Soc. B 282, 20151546 (2015).

17 Wassmann, P. et al. The contiguous domains of Arctic Ocean advection: Trails of life and death. Progress in Oceanography 139, 42–65 (2015).

18 Oziel, L. et al. Faster Atlantic currents drive poleward expansion of temperate phytoplankton in the Arctic Ocean. Nature communications 11, 1705 (2020).

19 Alabia, I. D., Garcia Molinos, J., Hirata, T., Mueter, F. J. & David, C. L. Pan-Arctic marine biodiversity and species co-occurrence patterns under recent climate. Scientific reports 13, 4076 (2023).

20 Polyakov, I. V. et al. Borealization of the Arctic Ocean in Response to Anomalous Advection From Sub-Arctic Seas. Front Mar Sci 7, 491 (2020).

21 Zehr, J. P. & Capone, D. G. Changing perspectives in marine nitrogen fixation. Science 368, eaay9514 (2020).

22 Karl, D. et al. The role of nitrogen fixation in biogeochemical cycling in the subtropical North Pacific Ocean. Nature 388, 533–538 (1997).

23 Capone, D. G. et al. Nitrogen fixation byTrichodesmiumspp.: An important source of new nitrogen to the tropical and subtropical North Atlantic Ocean. Global Biogeochemical Cycles 19, GB2024 (2005).

24 Shiozaki, T. et al. Basin scale variability of active diazotrophs and nitrogen fixation in the North Pacific, from the tropics to the subarctic Bering Sea. Global Biogeochemical Cycles 31, 996–1009 (2017).

25 Blais, M. et al. Nitrogen fixation and identification of potential diazotrophs in the Canadian Arctic. Global Biogeochemical Cycles 26, GB3022 (2012).

26 Harding, K. et al. Symbiotic unicellular cyanobacteria fix nitrogen in the Arctic Ocean. Proceedings of the National Academy of Sciences of the United States of America 115, 13371–13375 (2018).

27 Shiozaki, T. et al. Diazotroph community structure and the role of nitrogen fixation in the nitrogen cycle in the Chukchi Sea (western Arctic Ocean). Limnology and Oceanography 63, 2191–2205 (2018).

28 Shiozaki, T. et al. Biological nitrogen fixation detected under Antarctic sea ice. Nature Geoscience 13, 729–732 (2020).

29 Dugdale, R. C. & Goering, J. J. Uptake of New and Regenerated Forms of Nitrogen in Primary Productivity. Limnology and Oceanography 12, 196–206 (1967).

30 Yool, A., Martin, A. P., Fernandez, C. & Clark, D. R. The significance of nitrification for oceanic new production. Nature 447, 999–1002 (2007).

31 Fukai, Y. et al. Characteristics of autumn phytoplankton communities in the Chukchi Sea: resuspension of settled diatoms diatoms to the surface during strong wind events. Marine Ecology Progress Series 752, 35–50 (2025).

32 Fouilland, E., Gosselin, M., Rivkin, R. B., Vasseur, C. & Mostajir, B. Nitrogen uptake by heterotrophic bacteria and phytoplankton in Arctic surface waters. Journal of Plankton Research 29, 369–376 (2007).

33 Shiozaki, T. et al. Factors Regulating Nitrification in the Arctic Ocean: Potential Impact of Sea Ice Reduction and Ocean Acidification. Global Biogeochemical Cycles 33, 1085–1099 (2019).

34 Raimbault, P. & Garcia, N. Evidence for efficient regenerated production and dinitrogen fixation in nitrogen-deficient waters of the South Pacific Ocean: impact on new and export production estimates. Biogeosciences 5, 323–338 (2008).

35 Brown, S. M. & Jenkins, B. D. Profiling gene expression to distinguish the likely active diazotrophs from a sea of genetic potential in marine sediments. Environmental microbiology 16, 3128–3142 (2014).

36 Turk-Kubo, K. A. et al. Non-cyanobacterial diazotrophs: Global diversity, distribution, ecophysiology, and activity in marine waters. FEMS microbiology reviews 47, fuac046 (2023).

37 Shiozaki, T. et al. Distribution and survival strategies of endemic and cosmopolitan diazotrophs in the Arctic Ocean. The ISME journal 17, 1340–1350 (2023).

38 Coale, T. H. et al. Nitrogen-fixing organelle in a marine alga. Science 384, 217–222 (2024).

39 Pickart, R. S. et al. The Pacific water flow branches in the eastern Chukchi Sea. Progress in Oceanography 219, 103169 (2023).

40 Chen, Y. L. L., Chen, H. Y., Tuo, S. H. & Ohki, K. Seasonal dynamics of new production from Trichodesmium N-2 fixation and nitrate uptake in the upstream Kuroshio and South China Sea basin. Limnology and Oceanography 53, 1705–1721 (2008).

41 McLaughlin, F. A. & Carmack, E. C. Deepening of the nutricline and chlorophyll maximum in the Canada Basin interior, 2003-2009. Geophysical Research Letters 37, L24602 (2010).

42 Nishino, S., Itoh, M., Williams, W. J. & Semiletov, I. Shoaling of the nutricline with an increase in near-freezing temperature water in the Makarov Basin. Journal of Geophysical Research: Oceans 118, 635–649 (2013).

43 Nishino, S. et al. Atlantic-origin water extension into the Pacific Arctic induced an anomalous biogeochemical event. Nature communications 14, 6235 (2023).

44 Itoh, M., Nishino, S., Kawaguchi, Y. & Kikuchi, T. Barrow Canyon volume, heat, and freshwater fluxes revealed by long-term mooring observations between 2000 and 2008. J Geophys Res-Oceans 118, 4363–4379 (2013).

45 Wang, Y. et al. Enhanced wind mixing and deepened mixed layer in the Pacific Arctic shelf seas with low summer sea ice. Nature communications 15, 10389 (2024).

46 Cavalieri, D. J. & Parkinson, C. L. Arctic sea ice variability and trends, 1979–2010. The Cryosphere 6, 881–889 (2012).

47 Ingvaldsen, R. B. et al. Physical manifestations and ecological implications of Arctic Atlantification. Nature Reviews Earth & Environment 2, 874–889 (2021).

48 Polyakov, I. V. et al. Greater role for Atlantic inflows on sea-ice loss in the Eurasian Basin of the Arctic Ocean. Science 356, 285–291 (2017).

49 Juranek, L. W. Changing Biogeochemistry of the Arctic Ocean. Oceanography 35, 144–155 (2022).

50 Wu, J. F., Sunda, W., Boyle, E. A. & Karl, D. M. Phosphate depletion in the western North Atlantic Ocean. Science 289, 759–762 (2000).

51 Moore, C. M. et al. Large-scale distribution of Atlantic nitrogen fixation controlled by iron availability. Nature Geoscience 2, 867–871 (2009).

52 Mohr, W., Grosskopf, T., Wallace, D. W. R. & LaRoche, J. Methodological Underestimation of Oceanic Nitrogen Fixation Rates. PloS one 5, e12583 (2010).

53 Shiozaki, T. et al. Why is *Trichodesmium* abundant in the Kuroshio? Biogeosciences 12, 6931–6943 (2015).

54 Gradoville, M. R. et al. Diversity and activity of nitrogen-fixing communities across ocean basins. Limnology and Oceanography 62, 1895–1909 (2017).

55 Montoya, J. P., Voss, M., Kahler, P. & Capone, D. G. A simple, high-precision, high-sensitivity tracer assay for N-2 fixation. Applied and environmental microbiology 62, 986–993 (1996).

56 Kanda, J., Itoh, T., Ishikawa, D. & Watanabe, Y. Environmental control of nitrate uptake in the East China Sea. Deep-Sea Res Pt Ii 50, 403–422 (2003).

57 Shiozaki, T., Furuya, K., Kodama, T. & Takeda, S. Contribution of N2 fixation to new production in the western North Pacific Ocean along 155°E. Marine Ecology Progress Series 377, 19–32 (2009).

58 Shiozaki, T. et al. Nitrification and its influence on biogeochemical cycles from the equatorial Pacific to the Arctic Ocean. The ISME journal 10, 2184–2197 (2016).

59 Ward, B. B. & O’Mullan, G. D. Community Level Analysis: Genetic and Biogeochemical Approaches to Investigate Community Composition and Function in Aerobic Ammonia Oxidation. 397, 395–413 (2005).

60 Sigman, D. M. et al. A bacterial method for the nitrogen isotopic analysis of nitrate in seawater and freshwater. Analytical chemistry 73, 4145–4153 (2001).

61 Kawagucci, S. et al. Hadal water biogeochemistry over the Izu-Ogasawara Trench observed with a full-depth CTD-CMS. Ocean Science 14, 575–588 (2018).

62 Bolyen, E. et al. Reproducible, interactive, scalable and extensible microbiome data science using QIIME 2. Nature biotechnology 37, 852–857 (2019).

63 Martin, M. Cutadapt removes adapter sequences from high-throughput sequencing reads. EMBnet.journal 17, 10–12 (2011).

64 Callahan, B. J. et al. DADA2: High-resolution sample inference from Illumina amplicon data. Nature methods 13, 581–583 (2016).

65 Thompson, A. et al. Genetic diversity of the unicellular nitrogen-fixing cyanobacteria UCYN-A and its prymnesiophyte host. Environmental microbiology 16, 3238–3249 (2014).

66 Hasumi, H. CCSR ocean component model (COCO) version 4.0. (The University of Tokyo, 2006).

67 Watanabe, E., Onodera, J., Itoh, M. & Mizobata, K. Transport Processes of Seafloor Sediment From the Chukchi Shelf to the Western Arctic Basin. Journal of Geophysical Research: Oceans 127, e2021JC017958 (2022).

68 Reynolds, R. W., Rayner, N. A., Smith, T. M., Stokes, D. C. & Wang, W. An improved in situ and satellite SST analysis for climate. Journal of Climate 15, 1609–1625 (2002).

69 Woodgate, R. A., Aagaard, K., Swift, J. H., Smethie, W. M. & Falkner, K. K. Chukchi Borderland Cruise CBL 2002, Arctic West - Phase II (AWS-02-II). 51 pp (University of Washington, 2002).

70 Stemiletov, I. & Gustafsson, Ö. East Siberian Shelf Study Alleviates Scarcity of Observations. EOS 90, 145–146 (2009).

