## Supplementary Information for "Enhancing role of nitrogen fixation in biogeochemical cycles of the Pacific Arctic"

**This PDF file includes:**

Table S1

Figs. S1 to S5

**Other Supplementary Materials for this manuscript include the following:**

Database S1 to S2

Table S1 Pearson’s correlation matrix among environmental parameters and biological productivity in each year.

| **2015** | PAR | Temperature | Salinity | Nitrate | Ammonium | Phosphate | Chl *a* | N_2_ fix | PP | Nitrate assim | Nitrification |
| --- | --- | --- | --- | --- | --- | --- | --- | --- | --- | --- | --- |
| PAR | 1 |  |  |  |  |  |  |  |  |  |  |
| Temperature | 0.704^***^ | 1 |  |  |  |  |  |  |  |  |  |
| Salinity | 0.511^*^ | 0.826^***^ | 1 |  |  |  |  |  |  |  |  |
| Nitrate | -0.056 | 0.306 | 0.503^*^ | 1 |  |  |  |  |  |  |  |
| Ammonium | 0.013 | 0.476^*^ | 0.603^**^ | 0.941^***^ | 1 |  |  |  |  |  |  |
| Phosphate | 0.012 | 0.409 | 0.595^**^ | 0.984^***^ | 0.977^***^ | 1 |  |  |  |  |  |
| Chl *a* | -0.639^***^ | 0.809^***^ | 0.691^***^ | 0.056 | 0.232 | 0.143 | 1 |  |  |  |  |
| N_2_ fix | -0.337 | -0.330 | -0.223 | 0.236 | 0.113 | 0.172 | -0.258 | 1 |  |  |  |
| PP | 0.577^***^ | 0.722^***^ | 0.558^**^ | 0.199 | 0.293 | 0.227 | 0.885^***^ | -0.061 | 1 |  |  |
| Nitrate assim | 0.353 | 0.499^*^ | 0.584^**^ | 0.128 | 0.233 | 0.163 | 0.789^***^ | -0.248 | 0.741^***^ | 1 |  |
| Nitrification | -0.582^***^ | -0.214 | -0.008 | 0.472^*^ | 0.440^*^ | 0.433 | -0.216 | 0.395 | -0.140 | -0.048 | 1 |

| **2016** | PAR | Temperature | Salinity | Nitrate | Ammonium | Phosphate | Chl *a* | N_2_ fix | PP | Nitrate assim | Nitrification |
| --- | --- | --- | --- | --- | --- | --- | --- | --- | --- | --- | --- |
| PAR | 1 |  |  |  |  |  |  |  |  |  |  |
| Temperature | 0.538 | 1 |  |  |  |  |  |  |  |  |  |
| Salinity | 0.504 | 0.834^**^ | 1 |  |  |  |  |  |  |  |  |
| Nitrate | -0.792^**^ | -0.466 | -0.448 | 1 |  |  |  |  |  |  |  |
| Ammonium | -0.556^*^ | -0.376 | -0.644^*^ | 0.307 | 1 |  |  |  |  |  |  |
| Phosphate | -0.724^**^ | -0.437 | -0.636^*^ | 0.588^*^ | 0.509 | 1 |  |  |  |  |  |
| Chl *a* | 0.059 | 0.498 | 0.580^*^ | -0.075 | -0.222 | -0.025 | 1 |  |  |  |  |
| N_2_ fix | 0.367 | 0.502 | 0.437 | -0.484 | -0.459 | -0.329 | -0.185 | 1 |  |  |  |
| PP | 0.521 | 0.598^*^ | 0.659^*^ | -0.324 | -0.479 | -0.439 | 0.759^**^ | -0.045 | 1 |  |  |
| Nitrate assim | 0.415 | 0.498 | 0.514 | -0.175 | -0.434 | -0.300 | 0.706^**^ | -0.072 | 0.963^**^ | 1 |  |
| Nitrification | -0.313 | -0.410 | -0.545 | 0.182 | 0.280 | 0.475 | -0.325 | 0.189 | -0.404 | -0.201 | 1 |

| **2017** | PAR | Temperature | Salinity | Nitrate | Ammonium | Phosphate | Chl *a* | N_2_ fix | PP | Nitrate assim | Nitrification |
| --- | --- | --- | --- | --- | --- | --- | --- | --- | --- | --- | --- |
| PAR | 1 |  |  |  |  |  |  |  |  |  |  |
| Temperature | 0.504 | 1 |  |  |  |  |  |  |  |  |  |
| Salinity | 0.684^*^ | 0.685^*^ | 1 |  |  |  |  |  |  |  |  |
| Nitrate | 0.752^*^ | 0.324 | 0.551 | 1 |  |  |  |  |  |  |  |
| Ammonium | 0.584^*^ | 0.254 | 0.478 | 0.961^**^ | 1 |  |  |  |  |  |  |
| Phosphate | 0.726^*^ | 0.178 | 0.477 | 0.980^**^ | 0.944^**^ | 1 |  |  |  |  |  |
| Chl *a* | 0.838^**^ | 0.475 | 0.537 | 0.468 | 0.214 | 0.420 | 1 |  |  |  |  |
| N_2_ fix | -0.089 | 0.033 | 0.042 | -0.314 | -0.261 | -0.357 | -0.153 | 1 |  |  |  |
| PP | 0.857^**^ | 0.490 | 0.571 | 0.492 | 0.241 | 0.445 | 0.998^**^ | -0.149 | 1 |  |  |
| Nitrate assim | 0.777^*^ | 0.405 | 0.462 | 0.385 | 0.118 | 0.349 | 0.988^**^ | -0.180 | 0.985^**^ | 1 |  |
| Nitrification | -0.409 | -0.742 | -0.554 | -0.287 | -0.242 | -0.166 | -0.335 | 0.121 | -0.334 | -0.272 | 1 |

| **2020** | PAR | Temperature | Salinity | Nitrate | Ammonium | Phosphate | Chl *a* | N_2_ fix | PP | Nitrate assim | Nitrification |
| --- | --- | --- | --- | --- | --- | --- | --- | --- | --- | --- | --- |
| PAR | 1 |  |  |  |  |  |  |  |  |  |  |
| Temperature | 0.549 | 1 |  |  |  |  |  |  |  |  |  |
| Salinity | 0.192 | 0.757^*^ | 1 |  |  |  |  |  |  |  |  |
| Nitrate | 0.153 | 0.339 | 0.518 | 1 |  |  |  |  |  |  |  |
| Ammonium | 0.124 | 0.564^*^ | 0.706^*^ | 0.891^**^ | 1 |  |  |  |  |  |  |
| Phosphate | 0.196 | 0.517 | 0.693^*^ | 0.950^**^ | 0.974^**^ | 1 |  |  |  |  |  |
| Chl *a* | 0.807^**^ | 0.541 | 0.106 | -0.111 | -0.142 | -0.075 | 1 |  |  |  |  |
| N_2_ fix | n.a. | n.a. | n.a | n.a. | n.a. | n.a. | n.a. | 1 |  |  |  |
| PP | 0.703^*^ | 0.604^*^ | 0.165 | -0.101 | -0.145 | -0.087 | 0.933^**^ | n.a. | 1 |  |  |
| Nitrate assim | -0.257 | -0.143 | -0.021 | 0.467 | 0.177 | 0.260 | -0.126 | n.a. | 0.003 | 1 |  |
| Nitrification | -0.113 | -0.044 | 0.095 | 0.807^**^ | 0.526 | 0.612^*^ | -0.174 | n.a. | -0.122 | 0.851^**^ | 1 |

PAR, temperature, salinity, nitrate, ammonium, phosphate, and chlorophyll *a* (Chl *a*) were derived from surface data. Nitrogen fixation (N_2_ fix), primary production (PP), nitrate assimilation (Nitrate assim), and nitrification were derived from depth-integrated data.

^*^*p* < 0.05, ^**^*p* < 0.001

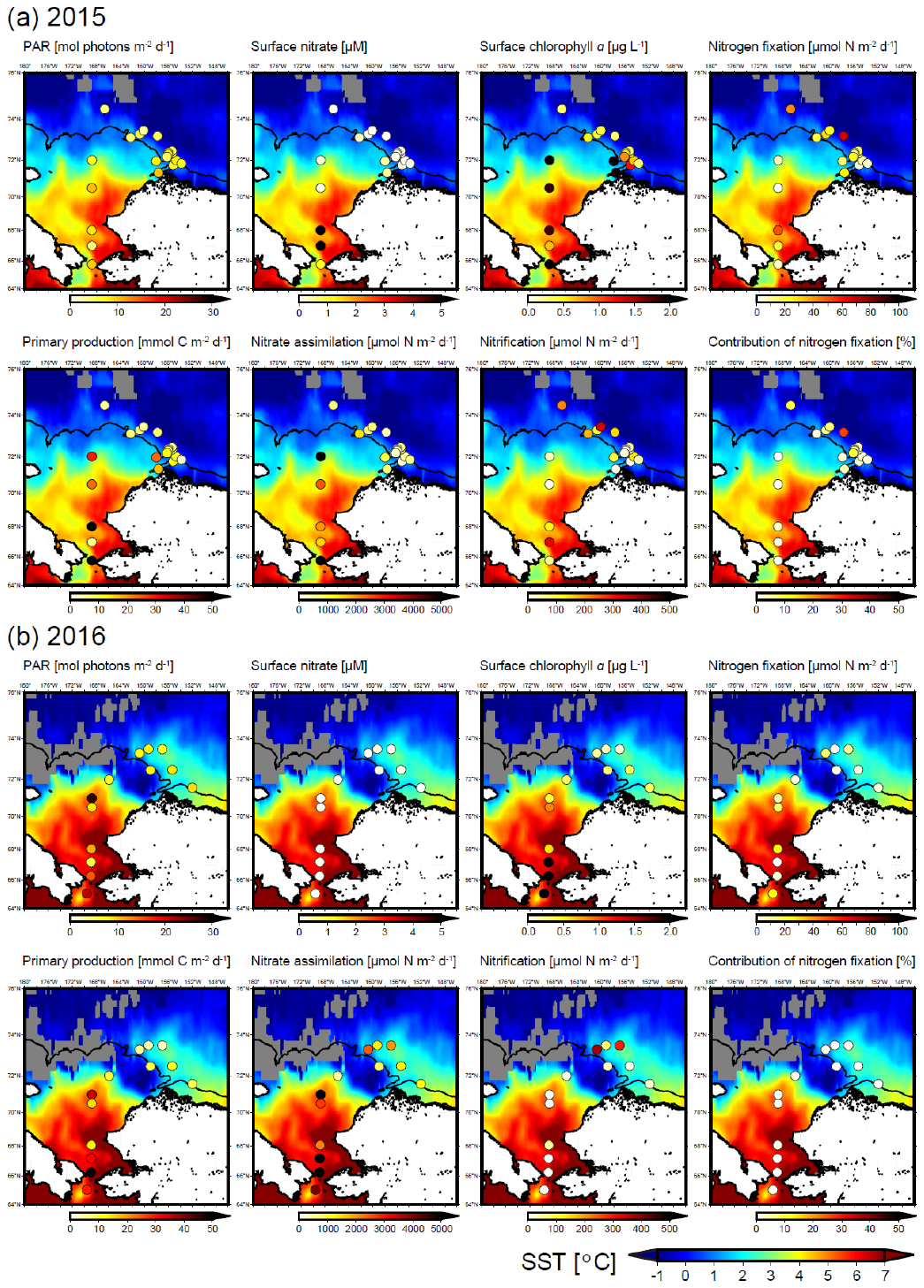

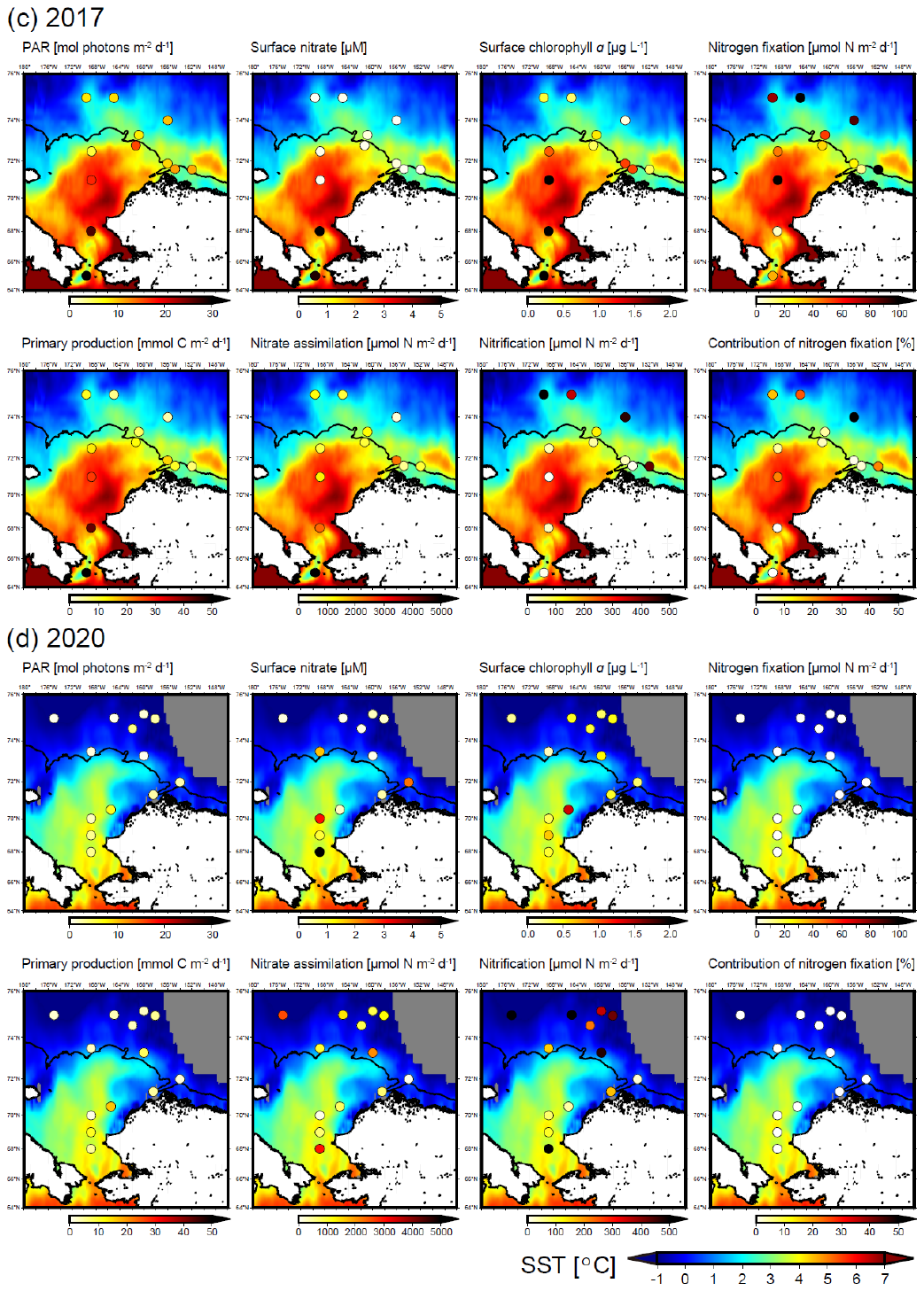

Fig. S1.

(a–d) Spatial distributions of PAR, surface nitrate, surface chlorophyll a, nitrogen fixation, primary production, nitrate assimilation, nitrification, and contribution of nitrogen fixation to new production in each year. Solid lines indicate 100-m isobaths.

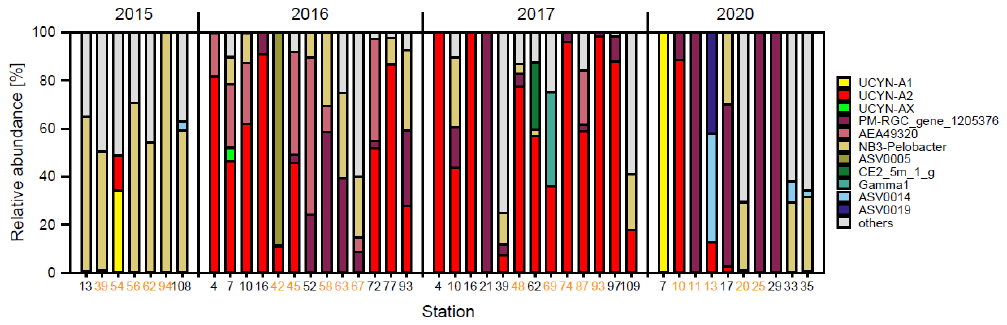

Fig. S2.

Diazotroph community structure in surface water in each year. Black and orange numbers indicate stations in shelf and off-shelf regions, respectively.

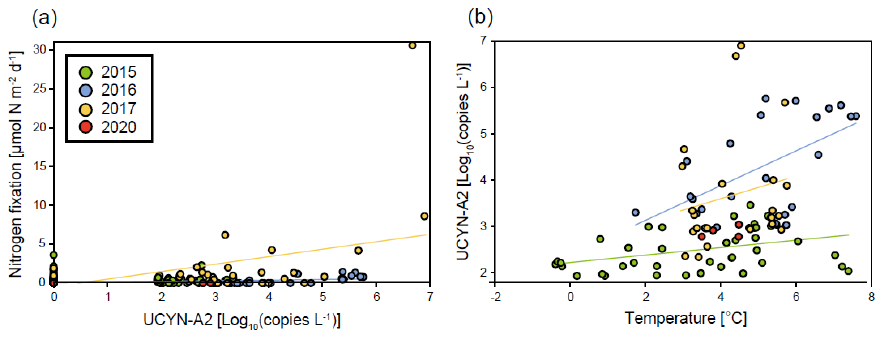

Fig. S3.

Relationships between (a) UCYN-A2 abundance and nitrogen fixation and (b) temperature and UCYN-A2 abundance in each year. Regression lines are plotted only for significant relationships (*P* < 0.05).

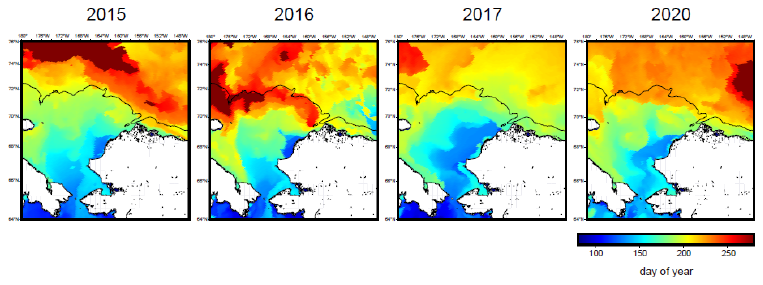

Fig. S4.

Timing of sea-ice retreat in each year.

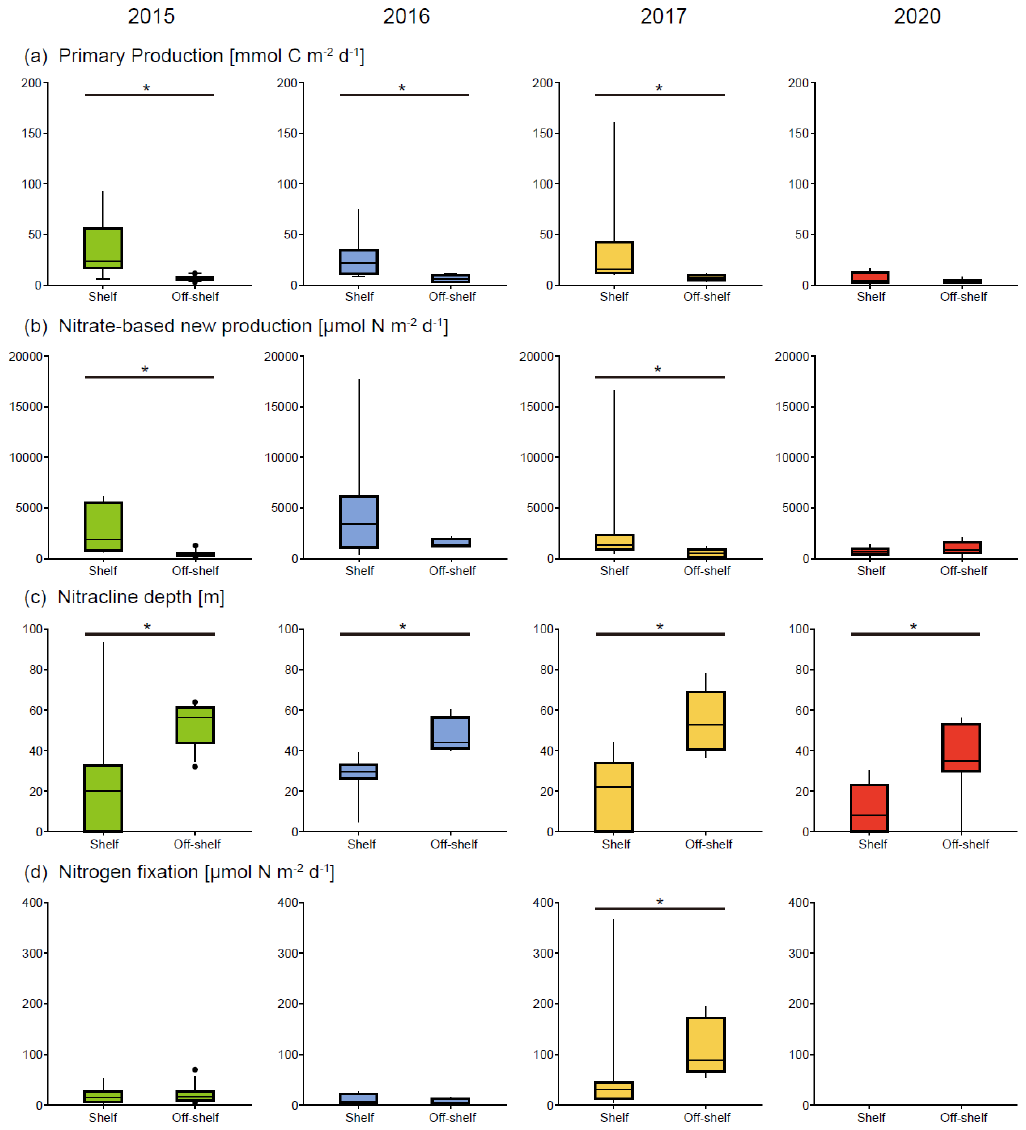

Fig. S5.

Differences in primary production, nitrate-based new production, nitracline depth, and nitrogen fixation between shelf and off-shelf regions. Lines on box plots indicate significant differences (*P* < 0.05, Wilcoxon rank sum test).

Dataset S1. (separate file)

Surface PAR, nitracline depth, depth-integrated biological activities, and contribution of N_2_ fixation to new production in each year.

Dataset S2. (separate file)

Cruise data at each light depth in each year.
